# Dynamic compensation, parameter identifiability, and equivariances

**DOI:** 10.1101/095828

**Authors:** Eduardo Sontag

## Abstract

A recent paper by Karin, Swisa, Glaser, Dor, and Alon introduced the mathematical notion of dynamical compensation (DC) in biological circuits, arguing that DC helps explain important features of glucose homeostasis as well as other key physiological regulatory mechanisms. The present paper establishes a connection between DC and two well-known notions in systems biology: system equivalence and parameter (un)identifiability. This recasting leads to effective tests for verifying DC in mathematical models.

## 1 Introduction

The recent paper [1] argued that physiological control systems should ensure a precise dynamical response despite variations in certain parameters, lest pathological conditions arise. The authors highlighted the biological significance of this robustness property through the analysis of several biological systems, including models of plasma glucose response in the face of changes in insulin sensitivity, parathyroid hormone control of calcium homeostasis, and arterial oxygen regulation in response to hypoxia. They formally introduced the system property of *dynamical compensation (DC)* with respect to variations in a parameter *p*, meaning that the complete output dynamics is exactly the same, for any time dependent input, independently of the precise value of *p*. They went on to provide a sufficient condition for DC to hold, and applied this condition to verify that their physiological examples have the DC property.

In this paper, we frame the notion of DC in the context of two related notions in systems biology: equivariances and parameter identifiability. We provide a necessary and sufficient condition for DC in the language of equivariances and partial differential equations, for which the sufficient condition given in [1] becomes a particular case, as does the condition given in [2] for fold-change detection (FCD) or input symmetry invariance, a Weber-like law in psychophysics [3]. We also point out that DC is equivalent to (structural) non-identifiability, and this in turn leads to the formulation of another necessary and sufficient condition for DC, this one using Lie-theoretic (directional) derivatives of observables along the vector fields defining the system.

## 2 Systems and equivalence

We formally state definitions in the language of dynamical systems with inputs and outputs, the standard paradigm in control systems theory [4]:

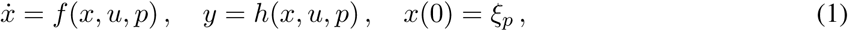

or, more explicitly,

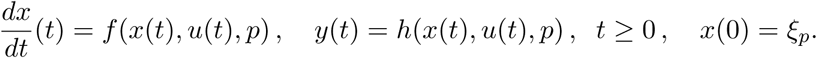

The functions *f*, *h* describe respectively the dynamics and the read-out map; *u* = *u*(*t*) is an input (stimulus, excitation) function, assumed to be piecewise continuous in time, *x*(*t*) is an *n*-dimensional vector of state variables, *y*(*t*) is the output (response, reporter) variable, and *p* is the parameter (or vector of parameters) that we wish to focus our attention on. In typical applications, *y*(*t*) = *x*_i_(*t*) is a coordinate of *x*. Values of states, inputs, outputs, and parameters are constrained to lie in particular subsets 𝕏, 𝕌, 𝕐, ℙ respectively, of Euclidean spaces ℝ*^n^*, ℝ*^m^*, ℝ*^q^*, ℝ*^s^*. Typically in biological applications, one picks 𝕏 as the set of positive vectors in ℝ*^n^*, *x* = (*x*_1_, *x*_2_,…, *x_n_*) such that *x_i_* > 0 for all *i*, and similarly for the remaining spaces. The state *ξ_p_* ∊ 𝕏 is an initial state, which could be different for different parameters; thus we view *ξ_p_* as a function ℙ → 𝕏. (We prefer the notation *ξ_p_* instead of *ξ*(*p*), so that we do not confuse *p* with the time variable.) For each input *u* : [0, ∞) → 𝕌, we write the solution of (1) with an initial condition *x*(0) =*ξ* (for example, *x*(0) =*ξ_p_*) as

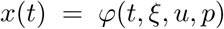

and the corresponding system response, which is obtained by evaluating the read-out map along trajectories, as

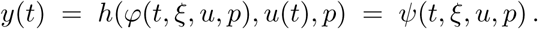

We assume that for each input and initial state (and each fixed parameter *p*), there is a unique solution of the initial-value problem *ẋ* = *f* (*x,u,p*), *x*(0) = *ξ* so that the mapping *φ* is well-defined; see [4] for more discussion, regularity of *f*, properties of ODE’s, global existence of solutions, etc. We refer to (1) as a *parametrized* (because of the explicit parameter) *initialized* (because of the specification of a given initial state for each parameter) family of systems.

Depending on the application, one might wish to impose additional restrictions on *ξ_p_*. For example, in [1] an additional requirement is that 0 ∊ 𝕌 and *ξ*_p_ ∊ 𝕏 is an equilibrium when *u*(0) = 0, which translates into *f* (*ξ_p_*, 0, *p*) = 0, and a similar requirement is made in [2] (this latter reference imposes the requirement that for each constant input *u*(*t*)≡μ there should exist an equilibrium, in fact). One may also impose stability requirements on this equilibrium; nothing changes in the results to be stated.

## 3 I/O Equivalence, Equivariances, and Identifiability

The question that we wish to address is as follows: provide conditions on the functions *f*, *h*, and *ξ* so that

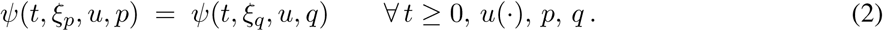

This is the “dynamic compensation” property from [1]. We prefer to use the terminology ℙ-*invariance*. This property says that the parameters *p* are (structurally) *unidentifiable* from input/output data [5,6].

In order to re-state this property in a more convenient form, let us fix from now on an arbitrary *p* ∊ ℙ, which we write as 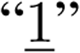 (for example, if ℙ is the set of all positive vectors (*p*_1_,…, *p*_s_), 1 could naturally be taken as the vector (1,1,…, 1)). Let us write 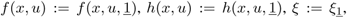 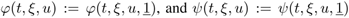. The following is a standard concept in control theory:

**Definition 1** *For any given parameter p, the system* (1) *is said to be* input/output *(I/O)* equivalent *to the system*

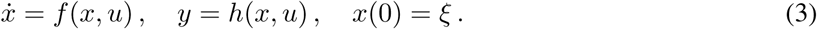

*provided, that*

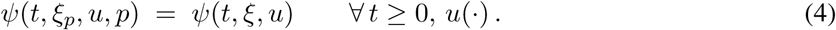

Obviously, (2) is equivalent to the property that, for every parameter *p*, (1) should be I/O equivalent to (3). Two approaches to testing I/O equivalence (or its lack) are described next.

### 3.1 Equivariances

**Definition 2** *Given a parametrized initialized family of systems (1), a set of differentiable mappings*

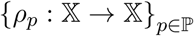

*is an* equivariance family *provided that:*

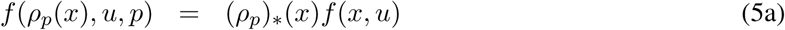

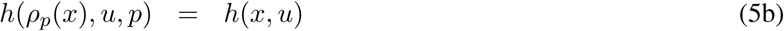

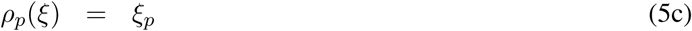

*for all x* ∊ 𝕏, *u* ∊ 𝕌, *and p* ∊ ℙ, *where in general p_*_ denotes the Jacobian matrix of a transformation ρ*. □ An important observation is as follows.

**Lemma 1** *If an equivariance family exists, then the parametrized family is ℙ-invariant.*

Observe that (5a) is a first order quasilinear partial differential equation on the components of the vector function *p_ρ_* (for each constant value *u* ∊ *U* and each parameter *p* ∊ ℙ), subject to the algebraic constraints given by (5b,c). Such equations are usually solved using the method of characteristics [7]. In principle, testing for existence of a solution through a “certificate” such as an equivariance is far simpler than testing all possible time-varying inputs *u*(*t*) in the definition of equivalence, in a fashion analogous to the use of Lyapunov functions for testing stability or of value functions in the Hamilton-Jacobi-Bellman formulation of optimal control theory [4]. The paper [2] discusses the relation to classical equivariances in actions of Lie groups and symmetry analysis of nonlinear dynamical systems.

**Proof of Lemma 1.** Fix a parameter *p* and an input *u*(·); we must show that (4) is satisfied. Let *x*(*t*) = *φ*(*t,ξ, u*), *t* ≥ 0, be the solution of the initial-value problem *ẋ = *f*(*x, u*),*x*(0) = ξ, so that *ψ*(*t, ξ, u*) = *h*(*x*(*t*),*u*(*t*)). Viewing the mapping *x* → *ρ_p_*(*x*) as a change of variables, we define *z*(*t*) := *ρ_p_*(*x*(*t*)).* Differentiating with respect to *t*,

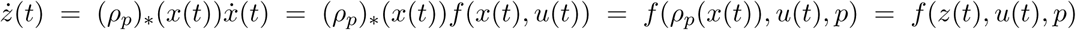

where we used (5a) applied with *x* = *x*(*t*) and *u* = *u*(*t*). Moreover, *z*(0) = *ρ_p_*(*x*(0)) = *ρ_p_*(*ξ*) =*ξ_p_*, because of (5c). Thus, since the solution of the initial-value problem *ẋ* = *f* (*x, u, p*), *x*(0) = *ξ_p_* is unique, it follows that *z*(*t*) = *φ*(*t,ξ_p_,u,p*). Now, by definition,

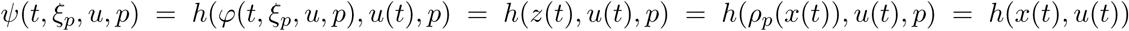

where the last equality follows from (5b). On the other hand, since *ψ*(*t,ξ,u*) = *h*(*x*(*t*), *u*(*t*)), we conclude that (4) holds, as desired.

It is a deeper result that the existence of an equivariance is also *necessary* as well as sufficient, see Section 4 for a statement and proof. Necessity tells us that it is always worth searching for an equivariance, when attempting to prove that a system has ℙ-invariance. Often one can guess such functions by looking at the structure of the equations.

### 3.2 Identifiability and Lie derivatives

For simplicity, in this section of the paper we restrict attention to systems in which inputs appear linearly; that is, systems defined by differential equations of the following general form:

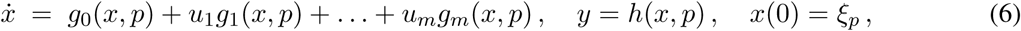

Linearity in inputs is not a serious restriction, nor is not having an explicit input in the read-out map, and it is easy to generalize to more general classes of systems, but the notations become far less elegant. We write the s coordinates of h as *h* = (*h*_1_,…, *h*_s_).

Consider the following set of functions (“elementary observables” of the system):

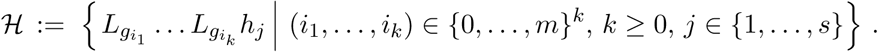

We are using the notation

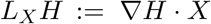

for the directional or Lie derivative of the function *H* with respect to the vector field *X*, and an expression as *L_Y_L_X_H* means an iteration *L_Y_*(*L_X_H*). We include in particular the case in which *k* = 0, in which case the expression in the defining formula is simply *h_j_*. For example, if the system is

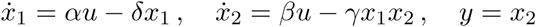

(initial states do not matter at this point; the parameters of interest could be all or a subset of *α, β, γ, δ*), then

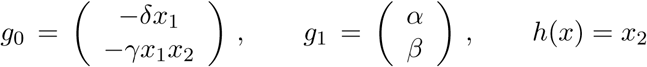

and we compute, for example,

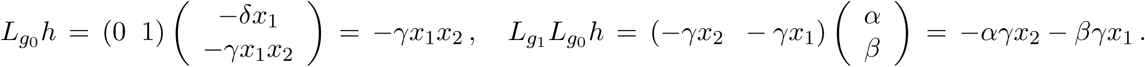

A necessary condition for invariance is as follows.

**Lemma 2** *If a system is* ℙ-*invariant, then all elements in* 𝓗, *when evaluated at the initial state, are independent of the values of parameters p.*

This condition is most useful when proving *non*-invariance. One simply finds one element of 𝓗 which depends on parameters. This test can also be used as a test for identifiability, that is to say the possibility of recovering all parameters perfectly from outputs: if one can reconstruct the values of *p* from the elementary observables (evaluated at the initial state), that is to say if the map

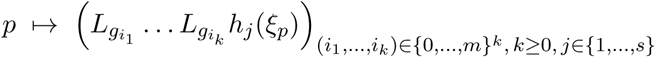

is one-to-one, then the parameters are identifiable. We discuss two examples in Section 3.4.

We sketch the proof of Lemma 2; see [4], Remark 6.4.2 for more details and references. Consider a piecewise constant control which is equal to *u*^1^ on [0, *t*_1_), equal to *u*^2^ on [*t*_1_,*t*_1_ + *t*_2_), …, and equal to *u^k^* on [*t*_1_ + … + *t*_*k*-1_, *t*_1_ + … + *t*_*k*_), starting from *x*(0) = *ξ_p_*. By invariance, the resulting output at time *t* = *t*_1_ + … + *t*_*k*_ is independent of the parameters. Let us denote the *j*th coordinate of this output value as

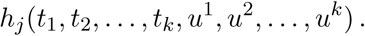

It follows that the derivatives with respect to the *t_i_*’s of this output are also independent of parameters, for every such piecewise constant control. By induction, one has

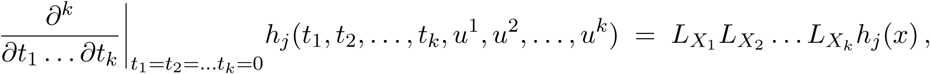

where we are using the notations 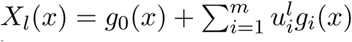 This expression is a multilinear function of the 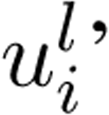, and a further derivation with respect to these control value coordinates gives the elementary observables, which therefore must also be independent of parameters.

We remark that, assuming a certain technical condition, a converse statement holds: if the elementary observables give the same values for some two parameters *p,q* ∊ ℙ, then the output for any possible inputs, whether step inputs or arbitrary inputs, is the same for these two parameters. The technical condition is that the vector fields *g_i_* are real-analytic, that is, that they can be expanded into locally convergent power series. The expressions that commonly appear in systems biology models, such as mass action kinetics (polynomial functions) or Hill functions (rational functions) are real analytic, as are trigonometric, logarithmic, and exponential functions, so analyticity is not a strong restriction on the theory. The proof of this converse fact consists two parts: (1) showing that outputs coincide for piecewise constant inputs, which is true because *h_j_* (*t*_1_,*t*_2_,…, *t_k_*, *u*^1^, *u*^2^,…, *u^k^*) can be expressed as a power series in terms of the observables, followed by (2) an approximation of arbitrary inputs by step ones (combining Lemma 2.8.2 with Proposition 6.1.11 in [4]; see also Section 4 in [8] and [9]).

### 3.3 Method of time derivatives at *t* = 0

A variation of the Lie derivatives technique for testing invariance (or lack thereof) is to employ inputs that are infinitely differentiable, checking if derivatives *y*(*t*), *y*′, *y*″ of outputs are independent of parameter values. For analytic systems, it suffices to check these derivatives at time *t* = 0 and to use analytic inputs (verified again by an approximation argument). Typically, all higher-order derivatives of outputs can be expressed as a function of a finite number of derivatives, which helps make computations effective using computer algebra packages. The theory is developed in [9-11], and is based on work by Fliess and others [12-16]. We discuss an example in Section 3.4.

### 3.4 Examples from the paper [1]

In its Supplement, the paper [1] presents the following class of systems that have ℙ-invariance, called there “dynamic compensation” (for arbitary *n*, but the main text specializes to *n* = 1):

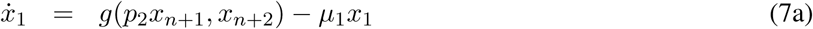

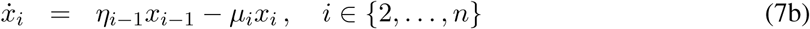

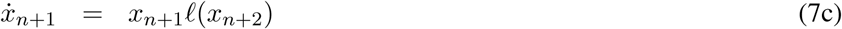

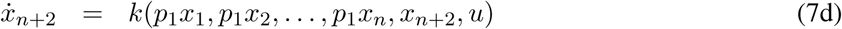

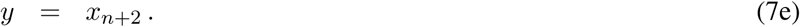

The state variables are positive, that is, 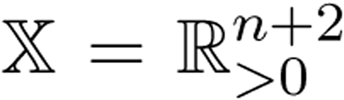, inputs and outputs are scalar and positive, 𝕌 = ℝ_>0_, 𝕐 = ℝ_>0_, and the parameters for which we desire invariance are two positive numbers, *p* = (*p*_1_,*p*_2_) ∊ ℙ = ℝ^2^_>o_. The additional parameters *η_i_* and *μ_i_* are fixed, and *g, k,l* are three scalar (differentiable) functions, with the following positive homogeneity property for *g:*

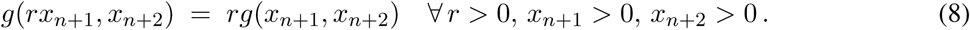

The initial state is implicitly specified in [1] by the requirement that for any parameter vector *p* = (*p*_1_, *p*_2_) there is a unique steady state when *u* = 0; we call this state *ξ_p_*. We take the reference parameter set to be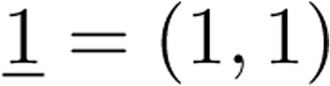.

In the paper [1], when *n* = 1 this system is motivated as a mechanism for a hormonal circuit in which the output *y* is a regulated variable, *x*_1_ is a hormone that regulates *y* = *x*_3_, and the variable *x*_n+1_ = *x*_2_ represents the functional mass of the tissue that secretes the hormone *x*_1_. There is feedback on the regulated variable *y*, with a gain *p*_1_, and a growth rate in *x*_n+1_ that is controlled by *y*, for instance through increase of proliferation. The parameter *p*_2_ quantifies the magnitude of the effect of the mass on the hormone production rate. In one of the examples in [1], the input *u*(*t*) is the meal intake of glucose, *x*_*n*+1_ represents *β* cells, *x*_1_ is insulin, and *y* is plasma glucose concentration; the coefficients *η_i_* correspond to transport rates, and the *μ_i_* combine degradation rate and transport rates. In the more general case *n* > 1, the coordinates *x*_1_,…, *x*_n_ represent the various physiological compartments that the hormone may circulate through.

To show ℙ-invariance, we exhibit an equivariance. In fact, the proof of dynamical compensation in [1] is based (without using that terminology) on showing that the following mapping is an equivariance family:

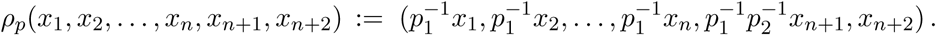

Since *h*(*x, u,p*) = *x*_*n*+2_ is not changed by *ρ*_p_, we have that *h*(*ρ_p_*(*x*), *u,p*) = *h*(*x, u*), and since as equivariances always map steady states into steady states, and steady states are assumed to be unique, there is no need to test the condition *ρ_p_*(*ξ*) = *ξ_p_*. So we only need to check that:

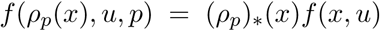

for all *p, x, u*. The map *ρ_ρ_* is linear, and its Jacobian matrix is diag 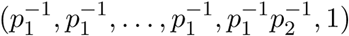, thus we need to verify:

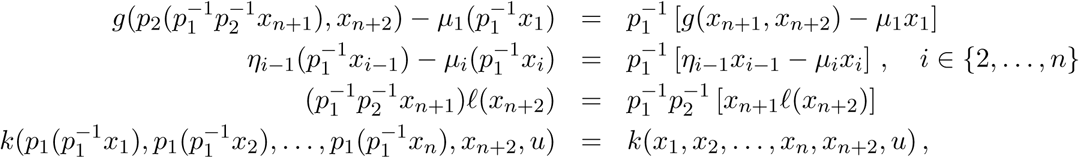

the last *n* + 1 of which are trivial, and the first one requires 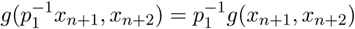, which holds because of the homogeneity property (8).

We next discuss the “linear integral feedback” and “linear proportional integral feedback” examples in the paper [1], which are given as examples of systems that are not 𝕡-invariant (the paper does this by simulation). The first system is:

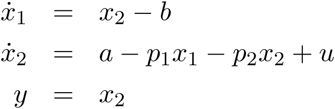

with the unique steady state*ξ_p_* = ((*a* − *p*_2_*b*)/*p*_1_, *b*) for *u*(0) = 0. Let us show that there is no possible equivariance *ρ_p_*(*x*_1_*x*_2_) = (*α_p_*(*x*_1_*x*_2_), *β_p_x*_1_*x*_2_). The requirement that *h*(*ρ_p_*(*x*),*u,p*) = *h*(*x,u*) means that *β_p_x*_1_,*x*_2_) = *x*_2_, so *ρ_p_*(*x*_1_,*x*_2_) = (*α_p_x*_1_*x*_2_)*x*_2_). The PDE *f*(*ρ_p_*(*x*),*u,p*)) = (*ρ_p_*)_*_(*x*)*f*(*x,u*) translates into:

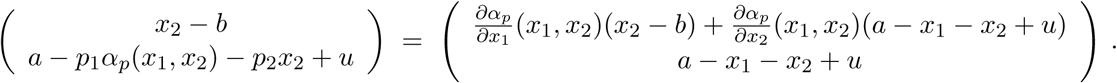

Comparing coefficients of *u* in the first component, we have that 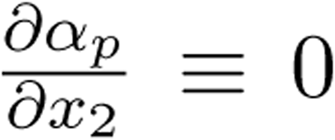, and this in turn implies that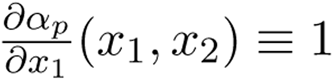. Thus the only possible choice is a linear function *α_p_*(*x*_1_,*x*_2_) = *c* + *x*_1_. On the other hand, comparing second components, we have

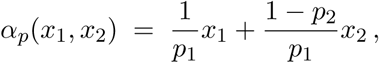

contradicting this formula. Thus, no possible equivariance exists, even if we only ask one of the parameters to vary. Combined with the necessity result to be shown below, this implies no invariance.

Another way to analyze this example is as follows. Let us compute the derivatives *d_i_* := *y^(i)^*(0^+^) when applying an input u(t) that is differentiable for t > 0. We have that *m_0_* = *b*, *m*_1_ = *u*_0_, *m*_2_ = −*p*_2_*u*_0_ + *u*_1_, 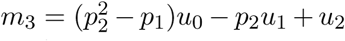, where *u* := *u*^(j)^ (0^+^). Therefore, from the output *y* (and in particular from its derivatives at time zero) we can reconstruct *p*_2_ and *p*_1_, for example using a step function *u* ≡ 1. This means that the parameter vector *p* is identifiable, which implies that the system is not ℙ-invariant.

Alternatively, using the formalism of Lie derivatives, with

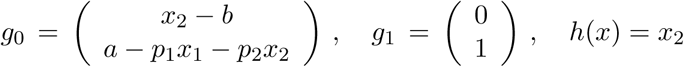

we can compute

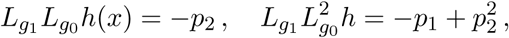

and we again see that we can recover the parameters, giving identifiability and hence non-invariance.

The second example of a system that fails invariance in the paper [1] is the “linear proportional integral feedback” given by:

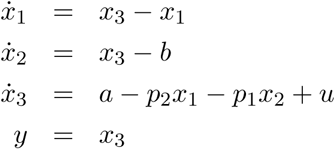

with the unique steady state *ξ_p_* = (*b*, (*a* − *p*_2_*b*)/*p*_1_,*b*) for *u*(0) = 0. The non-existence of equivariances is proved similarly to the previous example, and is not shown. For simplicity, we only perform the Lie derivative test. One can find

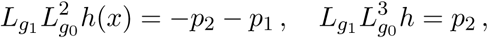

so once more we can recover the parameters, giving identifiability and hence non-invariance.

Let us now illustrate the method of derivatives with the example in Equation 7, which was already shown to be invariant by the method of equivariances. The equations are, in this case:

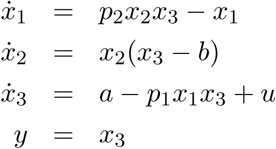

where *p* = (*p*_1_, *p*_2_) are the parameters of interest, *a, b* are other positive parameters, and the equations evolve in the positive orthant *x_i_* > 0. Using, for clarity, superscripts “(*i*)” to indicate derivative of order *i*, and since *y* = *x*_3_, we have that:

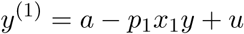

and thus, taking derivatives of this expression and substituting the equation for *ẋ* _1_:

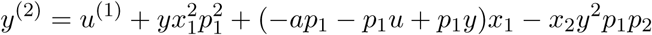

so, taking again derivatives and substituting the equations for *ẋ_1_* and *ẋ*_2_:

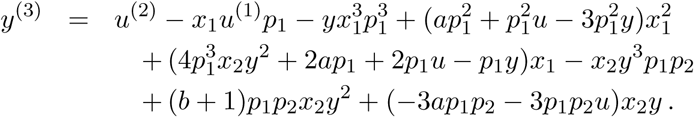

Observe that these are all functions of *t* (argument not shown). We can solve for *x*_1_ and *x*_2_ in terms of *y*^(1)^ and *y*^(2)^, obtaining:

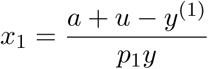

and

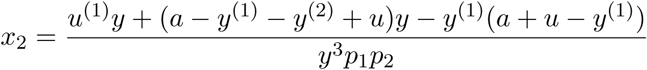

and, finally, substitute these expressions into the formula for *y*^(3)^, to obtain:

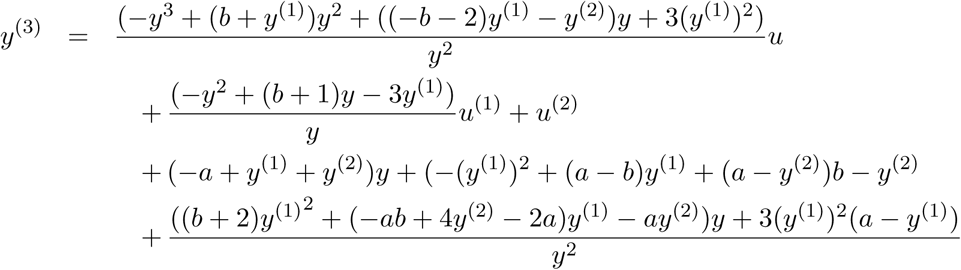

which involves only *y*, *y*^(1)^, and *y*^(2)^ but neither *x*_1_ and *x*_2_ nor the parameters *p*_1_ and *p*_2_. Taking derivatives on both sides, and iterating, this implies, in turn, that higher-order derivatives of *y* can also be expressed in terms of *y*, *y*^(1)^, and *y*^(2)^ and neither of the other state coordinates nor *p*_1_ and *p*_2_. This holds for all *t*, so in particular at *t* = 0. Therefore, if we prove that *y*(0), *y*^(1)^ (0), and *y*^(2)^ (0) are independent of parameters, the same will be true for all derivatives at *t* = 0, of any order, and therefore, because of analyticity (Taylor expansions and analytic continuation), it will follow that *y*(*t*) is independent of the parameters, for all *t* (and all inputs, even not differentiable, by the approximation argument mentioned earlier, combining Lemma 2.8.2 with Proposition 6.1.11 in [4]; see also Section 4 in [8] and [9]). Now, specializing at the equilibrium initial state

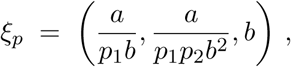

we have that, for any differentiable input, *y*(0) = *b*, *y*^(1)^(0) = *u*(0), and *y*^(2)^(0) = −(*a*/*b*)*u*(0) + *u*^(1)^(0). Thus, indeed neither of these derivatives depends on the parameters *p*, which confirms invariance.

It is interesting to observe that, in this example, there is no invariance to the other parameters, *a* and *b*; in fact, these parameters are identifiable. To see this, let us write, for convenience, *α* := *a*/*b*. Then,

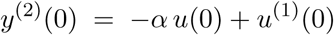

allows identification of *α*; for example, using the input *u*(*t*) ≡ 1, *Y*_2,1_ := *y*^(2)^(0) = −*α*, so can recover *α* = −*Y*_2,1_. From

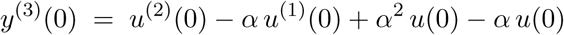

we cannot yet identify *a* and *b* individually. Taking one more derivative:

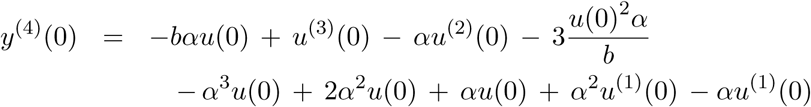

and therefore, if the input is *u*≡ 1, we obtain

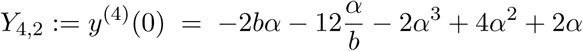

and if the input is *u* ≡ 2, we obtain

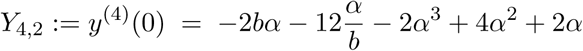

and combining these two results, we conclude that 2*Y*_4,1_ − *Y*_4,2_ = *6α*/*b*, allowing recovery of the parameter *b* (observe that *α* = *a*/*b* ≠ 0, since the parameters *a* and *b* are both positive).

### 3.5 Relations to FCD

The papers [2,3] studied a notion of scale invariance, or more generally invariance to input-field symmetries. We restrict attention here to linear scalings

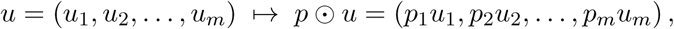

although the same ideas apply to more general symmetries as well. The property being studied is the invariance of the response of the following parametrized family of systems:

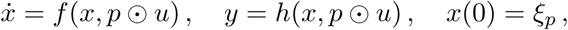

where the initial states *ξ_p_* are the (assumed unique) steady states associated to *p* ⊙ *u*(0). This property is also called “fold change detection” (FCD), because the only changes that are detectable are possibly “fold” changes in *u*, not simply scalings such as changes of units. It is clear that this is merely a special case of the current general setup.

## 4 Converses of the criteria

As pointed out earlier, the criterion in Lemma 2 admits a converse, provided that the system is defined by real-analytic functions: if all elements of 𝓗, when elaluated at *ξ_p_*, are independent of the values of the parameters *p*, then the system is ℙ-invariant. A similar result holds for identifiability: if one can recover all parameters as functions (perhaps nonlinear) of the elementary observables, then there is complete parameter identifiability (two different parameters give two different outputs, at least for some input).

The converse of Lemma 1 is also true, provided again that the maps *f* and *h* are real-analytic. An additional technical assumption, verified by all reasonable models including all examples in this paper, is that the systems are *irreducible,* meaning that it is accessible from the initial state, and observable. An accessible system is one for which the accessibility rank condition holds: 𝓕_LA_(*ξ*_p_) = ℝ^*n*^, where 𝓕_LA_ is the accessibility Lie algebra of the system. Intuitively, this means that no conservation laws restrict motions to proper submanifolds. For analytic systems, accessibility is equivalent to the property that the set of points reachable from has a nonempty interior; see a proof and more details in the textbook [4]. An observable system (for a fixed parameter *p* ξ ℙ, not shown in the notation) is one for which *ψ*(*t,ξ, u*) = *ψ*(*t, ξ′, u*) for all *u*, *t* implies *ξ* = *ξ*′. Intuitively, observability means that no pairs of distinct states can give rise to an identical temporal response to all possible inputs. For analytic input-affine systems, observability is equivalent to the property that any distinct two states can be separated by the observation space ([4], Remark 6.4.2). The proof of necessity is a corollary of a fundamental theorem in control theory, see the Appendix.

## A Appendix

We discuss here the notions of accessibility and observability required for the validity of a converse to Lemma 1, and give a proof of this converse result.

### A.1 Accessibility

We consider systems:

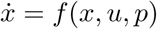

with the technical condition that *f* is a real-analytic functions of *x*. (Outputs do not matter for this section.) For any fixed parameter vector *p* ∊ ℙ, this system is said to be *accessible* from the initial state *ξ_p_* if the set of all states reached from *ξ_p_*,

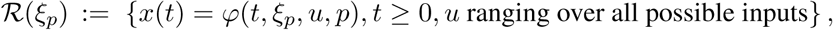

has a non-empty interior. In other words, 𝓡(*ξ_p_*) cannot be “too thin” (as would happen if there are conservation laws among the species *x_i_*, in which case one should first reduce the model to independent variables). See for instance the textbooks [4,17] for details. Accessibility can be tested as follows. The Lie bracket of two smooth vector functions 𝕏 → ℝ^n^ (more generally, one may extend to vector fields on a smooth manifold, but we restrict here to Euclidean spaces) is the function [*f, g*] : 𝕏 → ℝ^n^ defined by the following formula:

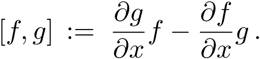

Let 𝓕 = {*f* (.,*u,p*),*u* ∊ 𝕌}, that is, the set of vector fields obtained when we plug-in an arbitrary constant input (still for our fixed parameter vector *p*). We now define the set 𝓕_∞_ that consists of all possible iterated [brackets *f_l_*…, [*f*_3_, [*f*_2_, *f*_1_]…]], over all choices of *f_i_*’s in 𝓕 and all *l* ≥ 1. Formally, 𝓕_∞_ is the union of the sets *F_k_* that are recursively defined as follows:

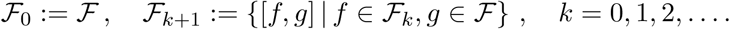

The *accessibility (or controllability) rank condition* is said to hold at *ξ_p_* if there are n (the dimension of the state space 𝕏) linearly independent vectors among all these vectors when evaluated at the initial state. An equivalent way to state this condition is in terms of the Lie algebra 𝓕_LA_ of vector fields generated by 𝓕, which is the linear span of 𝓕_∞_, the smallest Lie algebra of vector fields which contains 𝓕. The condition is that 𝓕_LA_(*ξ_p_*) = ℝ^n^. It can be shown that, for affine systems as in (6), 𝓕_LA_(*ξ_p_*) = {*g*_o_,…, *g_m_*}_LA_(*ξ_p_*).

As an illustration, take the example

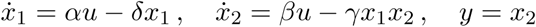

considered earlier. This is a system affine in inputs, with

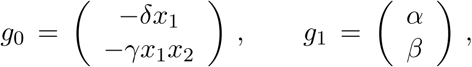

and we may compute:

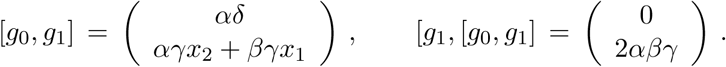

Since the determinant of the matrix with columns *g*_1_ and [*g*_1_, [*g*_0_, *g*_1_]] is equal to 2*α*^2^*βγ* ≠ 0 for every *x*, the accessibility rank condition holds, in particular, at the initial state.

### A.2 Observability

Consider again systems as in (1):

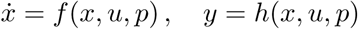

where *f* and *h* are real-analytic (initial states do not matter for this section). For any fixed parameter vector *p* ∊ ℙ, this system is said to be *observable* provided that, for any two distinct states *ξ* and *ξ*′, there is some input function *u* (which might depend on the particular pair of states *ξ* and *ξ*) such that the temporal responses *ψ*(*t,ξ,u*) and *ψ*(*t,ξ′, u*) differ at at least one time *t*. The equivalent contrapositive of this statement is “*ψ*(*t,ξ,u*) = *ψ*(*t,ξ′,u*) for all *u*, *t* implies *ξ* = *ξ*'.” For analytic input-affine systems (6) with output *h* = (*h*_1_,…, *h*_p_), one can restate the observability property as follows. We consider the set 𝓗 of elementary observables *L_gi_1__* …*L_gi_k__ h_j_*. Two states *ξ* and *ξ*′ are said to be *separated by* observables if there exists some *k* ∊ 𝓗 such that *k*(*ξ*) ≠ *k*(*ξ*′). Observability is equivalent to the property that any distinct two states can be separated by observables, or, in more intuitive terms, if all coordinates of the state can be obtained as (possibly nonlinear) functions of observables. This is analogous to the discussion of parameter identifiability, and its proof is also based on approximating arbitrary inputs by piecewise constant inputs, combining Lemma 2.8.2 and Proposition 6.1.11 in [4] (see also Section 4 in [8]).

As an illustration, take once more

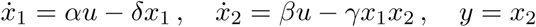

We need to show that the coordinates of the states can be recovered from the elements in 𝓗. Since *x*_2_ = *h*(*x*), so only need to give a formula for *x*_2_. Using

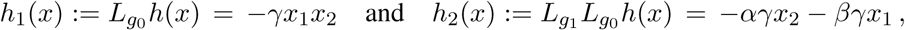

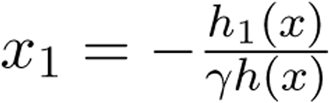, or, if we want to avoid division by zero when *y* = 0, 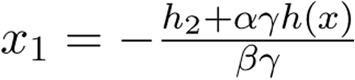.

### A.3 The converse to Lemma 1

Lemma 1 states that if an equivariance family exists, then the parametrized family is ℙ-invariant. We outline next how the converse of this Lemma follows from the theory of minimal realizations of nonlinear systems. Let us call a system *irreducible* if it is accessible from the initial state *ξ_p_*, and observable. We assume that the functions defining the system are analytic, and claim that, conversely, *if the parametrized family is* ℙ-*invariant and the systems are irreducible, then an equivariance family always exists.* To show this, let us fix any parameter *p* ∊ ℙ. We must find a differentiable mapping *ρ_p_* : 𝕏 → 𝕏 so that *ρ_p_*(*ξ*) = *ξ_p_* and *f* (*ρ_p_*(*x*),*u,p*) = (*ρ_p_*)*(*x*)*f*(*x,u*), *h*(*ρ_p_*(*x*),*u,p*) = *h*(*x,u*) for all *x* ∊ 𝕏 and *u* ∊ 𝕌, This means that *ρ* should be an isomorphism [18] between the two systems (1) and (3). Now, Theorem 5 in [18] shows that for two analytic and irreducible systems, I/O equivalence implies the existence of am isomorphism, which is exactly what we require.

